# Identification and predictability of soil quality factors and indicators from conventional soil and vegetation classifications

**DOI:** 10.1101/2021.03.04.433857

**Authors:** Paul Simfukwe, Paul W. Hill, Bridget A. Emmett, Davey L. Jones

**Affiliations:** Department of Natural Resources, School of Agriculture and Natural Resources, Mulungushi University, Great North Rd, Kabwe, Central Province, Zambia; School of Natural Sciences, Bangor University, Bangor, Gwynedd, LL57 2UW, UK; UK Centre for Ecology and Hydrology, Environment Centre Wales, Bangor, Gwynedd, LL57 2UW, UK

**Author notes:** Author Contributions: Conceived and designed the experiments: DLJ BAE PS. Performed the experiments: PS PWH. Analyzed the data: PS PWH DLJ Wrote the paper: PS PWH PS BAE DLJ.

**Keywords:** Soil health, Soil quality factor, Multivariate classification, Discriminant analysis, Cluster analysis

## Abstract

Generally, the physical, chemical and biological attributes of a soil combined with abiotic factors (e.g. climate and topography) drive pedogenesis. However, biological indicators of soil quality play no direct role in traditional soil classification and surveys. To support their inclusion in classification schemes, previous studies have shown that soil type is a key factor determining microbial community composition in arable soils. This suggests that soil type could be used as proxy for soil biological function and vice versa. In this study we assessed the relationship between soil biological indicators with either vegetation cover or soil type. A wide range of soil attributes were measured on soils from across the UK to investigate whether; (1) appropriate soil quality factors (SQFs) and indicators (SQIs) can be identified, (2) soil classification can predict SQIs; (3) which soil quality indicators were more effectively predicted by soil types, and (4) to what extent do soil types and/ or aggregate vegetation classes (AVCs) act as major regulators of SQIs. Factor analysis was used to group 20 soil attributes into six SQFs namely; *Soil organic matter*, *Organic matter humification*, *Soluble nitrogen*, *Microbial biomass*, *Reduced nitrogen* and *Soil humification index*. Of these, *Soil organic matter* was identified as the most important SQF in the discrimination of both soil types and AVCs. Among the measured soil attributes constituting the *Soil organic matter* factor were, microbial quotient and bulk density were the most important attributes for the discrimination of both individual soil types and AVCs. The *Soil organic matter* factor discriminated three soil type groupings and four aggregate vegetation class groupings. Only the Peat soil and Heath and bog AVC were distinctly discriminated from other groups. All other groups overlapped with one another, making it practically impossible to define reference values for each soil type or AVC. We conclude that conventionally classified soil types cannot predict the SQIs (or SQFs), but can be used in conjunction with the conventional soil classifications to characterise the soil types. The two-way ANOVA showed that the AVCs were a better regulator of the SQIs than the soil types and that they (AVCs) presented a significant effect on the soil type differences in the measured soil attributes.

## 1. Introduction

The multiple roles and functions of soil have resulted in several broad definitions of soil quality. One of the most widely adopted definitions for soil quality (SQ) was proposed by a committee for the Soil Science Society of America (chaired by Karlen) as: “the capacity of soil to function, within natural or managed ecosystem boundaries, to sustain plant and animal productivity, maintain or enhance water and air quality, and support human health and habitation” (Karlen et al., 1997). The quality of any soil has two parts: (1) the natural or inherent quality which is based on the parent geological material and soil-state-factors and is rather static, and (2) the dynamic soil quality which encompasses those soil properties that can change over relatively short time periods in response to human use and management (Carter, 2002; Fließbach et al., 2007; Bonfante et al., 2019). In contrast to the inherent SQ, the dynamic SQ can be used to monitor temporal trends on the same soil. There is no universally applicable set of inherent SQ criteria and optimum values (Carter, 2002) because soils with differences in the soil forming factors have different absolute capabilities (Seybold et al., 1998; Karlen et al., 2001). Therefore, soil quality and indicators have been defined by very different criteria and approaches dependent on the various functions the soil performs (Rapport et al., 1997; Carter, 2002, Cherubin et al., 2016). In spite of the lack of standard methodology and “critical limits”, it is possible to develop SQ ranges for specific soils evaluated with regard to specific land use and management regimes.

Soil quality is evaluated in terms of measurable soil attributes that measure specific physical, chemical, and biological properties; also known as soil quality indicators (SQIs; Shukla et al., 2006; Cherubin et al., 2016). Many of these properties are interrelated and the best SQIs are those that integrate and have the combined effect of several properties or processes that affect the capacity of a soil to perform a specified function (Dagnachew et al., 2019). SQIs should generally be linked and/or correlated with ecosystem processes and functions and should be responsive to variations in management and climate on an appropriate time scale (Doran and Safley, 1997, Bonfante et al., 2019). The SQIs which respond over the medium term i.e. those that are sensitive over few years and decades in land uses and management practices, may be the most useful for indicating soil quality changes as opposed to those which change either very rapidly (e.g. seasonally) or very slowly (e.g. over centuries) (Rapport et al., 1997; Dagnachew et al., 2019). Thus, measurement of key SQIs over time can be used to establish whether the quality of a soil under a given land use and management system is improving, declining or stable (Shukla et al., 2006; Ghaemi et al., 2014; Rayo et al.,2017).

Soil types are known to be inextricably determined by the physical, chemical and biological processes operating in soil, yet the biological indicators are rarely used in traditional soil classification and surveys (Cavigelli et al., 2005). Studies conducted by a number of researchers, such as Parkin (1993), Buyer et al. (2002), Girvan et al. (2003) and Ulrich and Becker (2006), have shown that soil type is a key factor determining microbial community composition in arable soils. Furthermore, Rapport et al. (1997) and Lagomarsino et al. (2009) reported that microorganisms and microbial communities can provide an integrated measure of soil quality; an aspect that cannot always be obtained with physical and chemical measures and/or analyses of higher organisms. Currently, bioindicators are mostly based on so-called sum or black-box parameters and generally include microbial indicators such as microbial biomass, activity and biodiversity (Rapport et al., 1997; Nielsen and Winding, 2002; Schloter, et al., 2018). Recently, an alternative has been proposed, based on the use of specific ratios that report on function such as the quotients of *microbial respiration-C-to-microbial biomass-C* (*q*CO_2_) and the *microbial biomass-C-to-organic matter-C* ratio (*q*Mic) (Schloter, et al., 2018). These indicators avoids the problems of comparing trends in soils with different organic matter or microbial biomass content and appears to provide a more sensitive indicator of soil changes than either activity or population measurements alone (Lagomarsino et al., 2009). In this study, we used multivariate statistical methods to explore these relationships using 20 physico-chemical and biological soil properties as Total Data Set (TDS). Using factor analysis the 20 correlated variables were reduced to 6 uncorrelated factors (soil quality factors; SQFs) also called Minimum Data Set (MDS) that were linear functions of the original 20 variables. The main questions addressed in this study were: (1) Can soil classification be used to predict SQFs and SQIs? (2) Which SQFs and SQIs are more effectively predictable by soil type in UK soils? (3) To what extent do soil types and/or AVCs act as major regulators of SQFs or SQIs?

## 2. Materials and methods

### 2.1. Soil sampling and preparation

Soil samples were collected throughout the UK as part of the Centre for Ecology and Hydrology Countryside Survey (CS) 2007 (Emmett et al., 2010) with sites representing the main types of landscape and soil groups. To encompass all the major soil and land use types, a total of 304 soil samples were collected throughout the UK, based on a stratified random sample of 1 km squares at gridpoints on a 15 km grid using the Institute of Terrestrial Ecology (ITE) Land Classification as the basis of the stratification (Scott, 2008). Figure S1 shows the general location and distribution of samples across the UK. At each grid intersection, a 1 km^2^ sample area was selected. Within the 1 km^2^ sample area, 3 plots (5 × 5 m^2^) were randomly located and a single 15 cm long × 4 cm diameter soil sample was collected from each of the plots. Topsoils were only selected for sampling to reflect standard practice in national monitoring schemes (Bellamy et al., 2005) such as Soil Survey England and Wales handbook (Hodgson, 1976), the National Soil Monitoring Network (Emmett, B.A., 2006) and the UK Soil Monitoring Network (Environmental Agency, 2008).

The soil leachate was collected according to Emmett et al. (2008). The soil leachate replicate cores were first wetted to field capacity with artificial rainfall (125 μM NaCl, 15.7 μM CaCl2, 1.3 μM CaSO4, 15.3 μM MgSO4, 12.3 μM H2SO4) in the dark at 10°C until the soils were fully wetted. The cores were then sprayed with artificial rainfall until a further 150 ml of leachate had been collected resulting in a solution with a pH of approximately 4.6. After washing out the cores, a small amount of suction was applied to drain larger pores. Cores were then incubated under anaerobic conditions for 4 weeks, at 10 °C, approximately UK mean summer soil temperature Cores were then extracted with 1 M KCl, and ammonium and nitrate concentrations were determined as a measurement of mineralisable N using a TOC-VCSH/CSN analyzer (Shimadzu Corp., Kyoto, Japan) as describe below.

Across all land uses (Supplementary Information S2) and aggregate vegetation class (AVC) categories, the dominant soil types (% of total) were: Brown soils (32%), Groundwater gleys (13%), Surface water gleys (19%), Lithomorphic soils (8 %), Peats (15%), Pelosols (2%) and Podzolic soils (11%). See Table S1 for their equivalents in the WRB classification. All the sites were characterised by a temperate climate with a North-South mean annual temperature range of 7.5 to 10.6°C and East-West mean annual rainfall range of 650 to 1700 mm.

**Table 1.** Summary of the Aggregate Vegetation Classes (AVCs) used for assessment of vegetation condition. The brackets indicate the abbreviation of the vegetation class (adapted from Smart et al., 2003).

| Aggregate vegetation class (AVC) +(abrev) | Description |
| --- | --- |
| 1. Crops and weeds (Craw) | Weedy communities of cultivated and disturbed ground, including species-poor arable and horticultural crops. |
| 2. Tall grass and herbs (Tgah) | Less intensively managed tall herbaceous vegetation typical of field edges, roadside verges, stream sides and hedge bottoms. |
| 3. Fertile grassland (Frtg) | Agriculturally improved or semi improved grassland. Often intensively managed agricultural swards with moderate to high abundance of perennial rye grass. |
| 4. Infertile grassland (Infg) | Less-productive, unimproved and often species rich grasslands in a wide range of wet to dry and acid to basic situations. |
| 5. Lowland wooded (Lwlw) | Vegetation dominated by shrubs and trees in neutral or basic situations, generally in lowland Britain. Includes many hedgerows. |
| 6. Upland wooded (Uplw) | Vegetation of broadleaved and conifer woodland often in more acidic situations, generally in upland Britain. |
| 7. Moorland grass mosaics (Mrgm) | Extensive, often unenclosed and sheep grazed hill pastures throughout Britain. |
| 8. Heath and bog (Htab) | Vegetation dominated by heathers. Includes drier heaths as well as bog. Mostly in the uplands. |

### 2.2. Aggregate vegetation classes

The vegetation data from the plots were analysed using the classification by Aggregate Classes (ACs) or Aggregate Vegetation Classes (AVCs). The AVCs were the vegetation types produced from a quantitative hierarchical classification of the different species found in sample plots. The eight AVCs used for assessing vegetation condition are listed in Table 1. Across all the soils sampled, the AVCs represented (% of the total): 18% Crop and weeds, 17% Fertile grasslands, 22% Heath and bogs, 20% Infertile grasslands, 2% Lowland woodland, 10% Moorland grass mosaics, 4% Tall grass and herbs and 7% Upland woodland.

### 2.3. Soil analysis

Soil pH was determined in soil-distilled water extracts (1:2.5 w/v soil to water soil ratio) using a glass electrode (Gelplas general purpose electrode, BDH) and HI-209 pH meter (Orion research, Boston, MA, USA). Soil moisture was determined by weight loss after oven drying at 105°C overnight (>16 h). Water content at field capacity was estimated by saturating the soil followed by measuring the water retained in the soil at −33 kPa. Bulk density was calculated (mass/volume) after removal of stones from the cores (>2 mm in diameter). Loss on ignition (LOI) was undertaken at 375°C for 16 h. Soil organic carbon (SOC) was calculated from the LOI values according to the method of Ball (1964) where

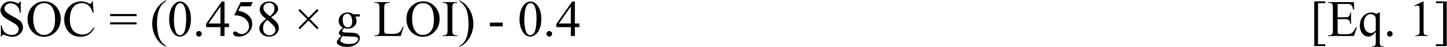

Phosphorus was determined by the Olsen P method according to Emmett et al. (2008). Total C and N were determined using UKAS accredited method SOP3102 on an Elementar Vario-EL elemental analyser (Elementaranalysensysteme GmbH, Hanau, Germany) according to Emmett et al. (2008, and 2010).

Soil respiration (SR) was determined on a 15 cm long, 2.5 cm diameter soil cores with a 1250 cm^3^ head space. The soils were incubated at 10°C for 1 h (at which linearity was established). Subsequently, the head space gas was analysed for CO_2_ concentration using a Clarus 500 Gas Chromatograph (Perkin Elmer Corp., Beverley, MA). The CO_2_ flux was established by comparing the CO_2_ concentration before and after incubation. Soil microbial biomass C and N were estimated on moist soil samples (10 g) using the modified chloroform-fumigation-extraction (CFE) method of Vance et al. (1987). For each soil 10g of the control and the fumigated samples were extracted with 1 M KCl. The TOC and TON in the 1 M KCl extracts was determined using a TOC-VCSH/CSN analyzer (Shimadzu Corp., Kyoto, Japan). Extraction efficiency correction factors of 0.45 and 0.54 were used for microbial C and N, respectively (Joergensen and Mueller, 1996a; 1996b; Fließbach et al., 2006). Soil microbial biomass was therefore calculated according to the formula: C_mic_ = EC/*k*EC, where EC = (TOC in fumigated samples - TOC in control samples) and *k*EC = 0.45, and N_mic_ = EN/*k*EN, where EN = (total N in fumigated samples – total N in control samples) and *k*EN = 0.54. The microbial C:N ratios were subsequently calculated from these values.

The metabolic and microbial quotients were calculated indices. The metabolic quotient or coefficient was calculated as the ratio between the CO_2_-C from basal respiration and the microbial biomass-C (CO_2_-C_resp_-to-C_mic_), expressed as µg CO_2_-C mg^-1^ biomass-C h^-1^. It is also known as the specific respiration rate (*q*CO_2_) (Anderson and Domsch, 1993). The microbial quotient was calculated as the ratio between the microbial biomass-C-to-total organic C (C_mic_-to-C_org_).

### 2.4. Leachate analysis

Leachate dissolved organic C (DOC) and total organic N (TON) were measured using a TOC-VCSH/CSN analyzer (Shimadzu Corp., Kyoto, Japan) and the DOC:TON ratio subsequently calculated. Nitrate and ammonium concentrations were measured with a Skalar SAN^++^ segmented-flow autoanalyser (Skalar, Breda, Netherlands), based on the cadmium (Cd) reduction method (Maynard and Kalra, 1993; Griffin, et al., 1995) and the modified Berthelot reaction (Searie, 1984) respectively. Electrical conductivity (EC) was measured with a standard platinum 1 cm electrode on a 4520-EC meter (Jenway Ltd, Dunmow, Essex, UK). pH was measured using a glass electrode (Gelplas general purpose electrode, BDH) on a HI-209 pH meter (Orion research, Boston, MA, USA). Total free amino acids were determined using the fluorometric OPAME procedure of Jones et al. (2002) and a Cary Eclipse Fluorescence Spectrophotometer (Varian Inc., Australia) using a leucine standard. Humic substances were determined by measuring the absorbance of 350 µl of leachate at 254 and 400 nm (UV and visible range respectively) on a PowerWave XS scanning microplate spectrophotometer (BioTek^®^ Instruments, Winooski, VT). The absorbance of deionised water was used as a control. A humification index (HIX) was calculated by dividing the absorbance at 254 nm by the absorbance at 400 nm (Zsolnay et al., 1999; Embachar et al., 2007). Soluble phenolic concentrations were assayed using a modification of the method of Box (1983) and Ohno and First (1998) using Na_2_CO_3_ (1.9 M) and the Folin-Ciocalteu reagent (Sigma-Aldrich, Poole, Dorset) (DeForest et al., 2005). The blue-coloured phenolics were measured at 750 nm using a PowerWave XS scanning microplate spectrophotometer (BioTek^®^ Instruments).

### 2.5. Statistical analyses

ANOVA, Factor, Discriminant and Cluster analyses were all determined using SPSS version 15.0 (SPSS Inc., Chicago, IL) and GenStat version 8 (VSN International Ltd, Hemel Hempstead, UK). They were used to analyse the measured attributes to investigate the effect of soil types and AVCs on the SQIs identified. To identify significant differences between treatments, post hoc multiple comparison (pair-wise) tests were made using the Gabriel test where homogeneity of variance was assumed and Games-Howell procedure where unequal variance occurred. Some variables were clearly not normally distributed judging from the Q-Q plots (data not presented); however, all the factors (SQFs) from factor analysis and discriminant analysis were normally distributed.

For the cluster analysis, the average linkage method and a squared Euclidean distance measure were used with a rescaled distance cluster combined measure on the similarity axis. The variables were standardized to minimize the effect of scale differences since the variables possessed different units.

## 3. Results

### 3.1. Biological, physical and chemical properties of soils

The variability of individual soil quality indicators across the range of soil types is shown in Figure 1. The box plots shows the spread of each measured soil property for each soil type as well as the data’s symmetry and skewness. (The boundary of the box closest to zero indicates the 25^th^ percentile, the line within the box marks the median (50^th^ percentile), and the boundary of the box farthest from zero indicates the 75^th^ percentile while the whiskers below and above the box indicate the 10^th^ and 90^th^ percentiles where outliers are present). From the box plots, most of the soil quality indicators did not show differentiations among the soil types save for the following: microbial quotient, SOC and Soil Respiration separated the peats from the rest; pH and C:N leachate separated the peats and the podzols from the rest, while the bulk density grouped the soils in three groups of Pelosols, the Browns, Ground-water gleys and the Surface-water gleys (average =1.1 g cm^-3^) in one group; Podzols and Lithomorphics (av = 0.5 g cm^-3^) in the second group and peats (ave 0.2 g cm^-3^) in the third group. All other properties were did not show effective differentiations among the soil types.

**Figure 1:**
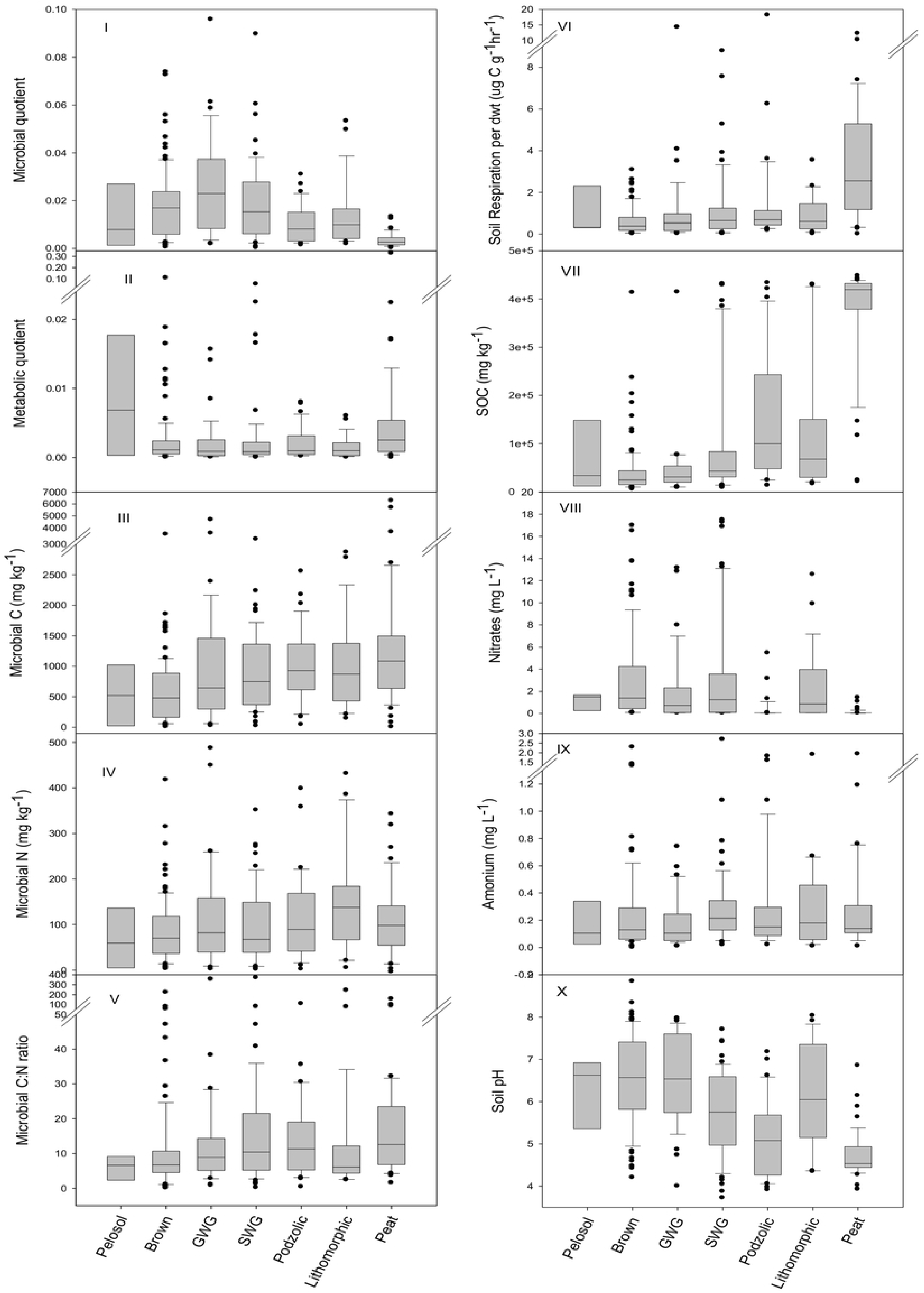
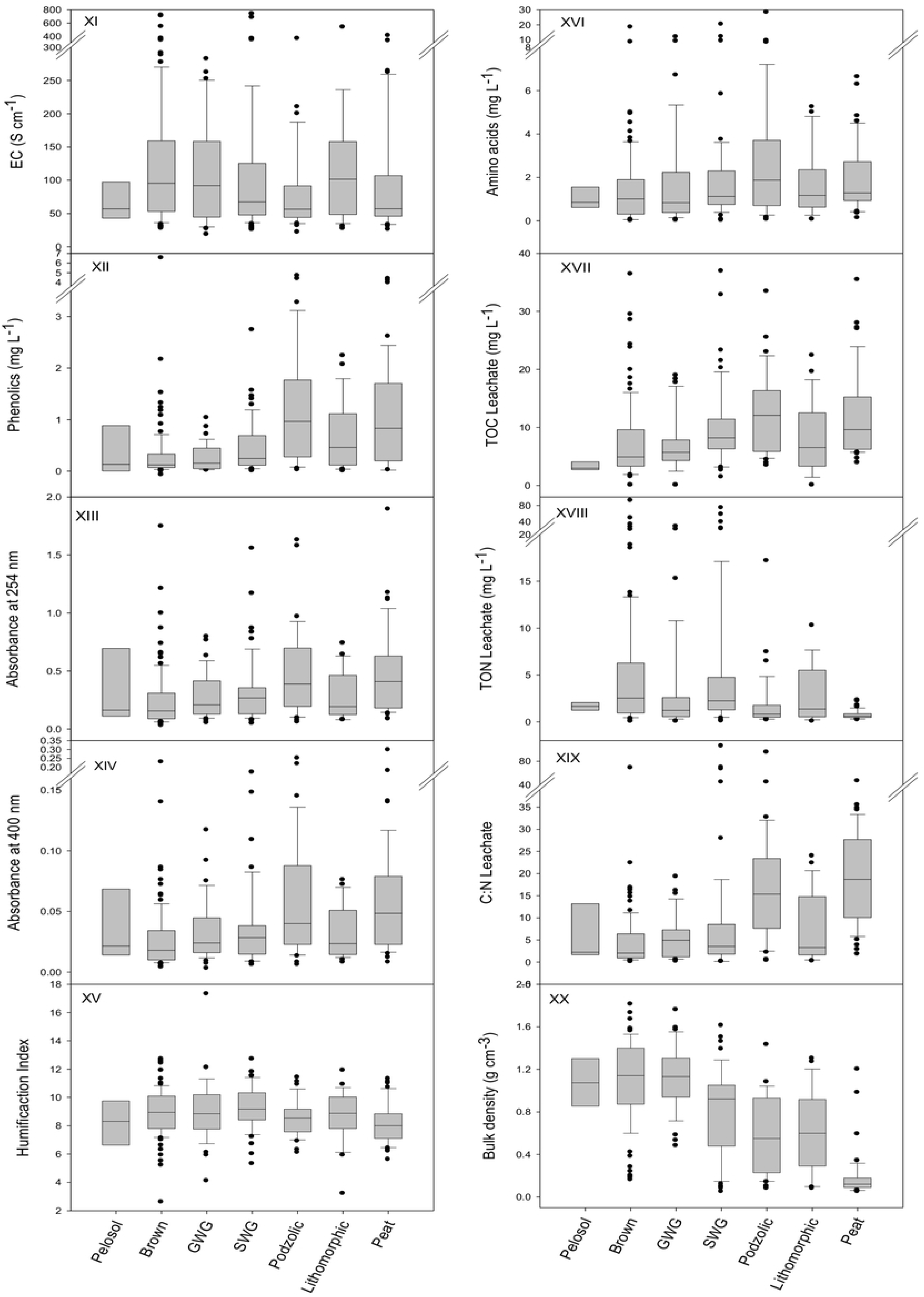
Box plots showing the spread of each measured soil property for each of the major soil types from 304 individual soils sampled as part of a nationwide soil quality assessment in UK. The boundary of the box closest to zero indicates the 25^th^ percentile, the line within the box marks the median (50^th^ percentile), and the boundary of the box farthest from zero indicates the 75^th^ percentile. Whiskers below and above the box indicate the 10^th^ and 90^th^ percentiles where outliers are present. GWG and SWG represent groundwater and surface water gley soils respectively.

### 3.2. Relationships among soil properties

Correlation analysis of the 20 soil attributes representing soil biological, physical and chemical properties resulted in significant correlation (*P <* 0.05) in 112 of the 190 soil attribute pairs (Table 2). Of these, the highest significant (*P* < 0.01) positive correlations was between humic substances at 254 nm versus those at 400 nm (*r* = 0.97). Other highly significant (*P* < 0.01) positive correlations were between the absorbance at 254 nm or 400 nm versus DOC (*r* = 0.78 and *r* = 0.71 respectively); leachate TON versus NO_3_ (*r* = 0.78), and bulk density versus pH (*r* = 0.70). Additional notable significant (*P* < 0.01) positive correlations (*r* > 0.50) were between: microbial-N versus microbial-C, SOC versus soil respiration, the leachate C:N ratio versus SOC, electrical conductivity versus both nitrate and TON, phenolics versus absorbance at 254 nm and DOC versus absorbance at 400 nm. The highest significant (*P* < 0.01) negative correlation was between bulk density versus SOC (*r* = −0.83) Other notable significant (*P* < 0.01) negative correlations were between: bulk density versus either microbial-C (*r* = −0.42), soil respiration (*r* = −0.51) or the leachate C:N ratio (*r* = −0.47); SOC versus *q*Mic (*r* = −0.47) and pH versus either SOC (*r* = −0.66), absorbance at 400 nm (*r* = −0.42), leachate DOC (*r* = −0.40) or leachate C:N ratio (*r* = −0.47)

**Table 2.** Correlations among physical, chemical and biological soil attributes

| Variable | qMic | qCO2 | Mic C | Mic N | Mic CN | SR | SOC | Nitrate | Amonia | pH |
| --- | --- | --- | --- | --- | --- | --- | --- | --- | --- | --- |
| qMic | 1 |  |  |  |  |  |  |  |  |  |
| qCO2 | -0.07 | 1 |  |  |  |  |  |  |  |  |
| Mic C | 0.23(**) | -0.07 | 1 |  |  |  |  |  |  |  |
| Mic N | 0.17(**) | -0.05 | 0.63(**) | 1 |  |  |  |  |  |  |
| Mic CN | 0.18(**) | -0.02 | 0.03 | -0.24(**) | 1 |  |  |  |  |  |
| SR | -0.26(**) | -0.01 | 0.31(**) | 0.09 | -0.02 | 1 |  |  |  |  |
| SOC | -0.47(**) | -0.04 | 0.39(**) | 0.09 | 0.04 | 0.61(**) | 1 |  |  |  |
| Nitrate | 0.21(**) | -0.01 | -0.15(**) | -0.12(*) | 0.20(**) | -0.22(**) | -0.33(**) | 1 |  |  |
| Amonia | -0.05 | -0.04 | 0.08 | 0.06 | -0.03 | 0.02 | 0.04 | 0.06 | 1 |  |
| pH | 0.35(**) | 0.08 | -0.31(**) | 0 | -0.11(*) | -0.39(**) | -0.66(**) | 0.25(**) | -0.18(**) | 1 |
| Ec | 0.03 | 0.06 | -0.09 | 0.01 | 0.17(**) | -0.08 | -0.03 | 0.59(**) | 0.03 | 0.12(*) |
| Phenols | -0.23(**) | -0.01 | 0.19(**) | 0.08 | -0.01 | 0.27(**) | 0.39(**) | -0.19(**) | 0.38(**) | -0.36(**) |
| Absb @ 254 | -0.24(**) | -0.01 | 0.10(*) | -0.03 | 0.05 | 0.22(**) | 0.34(**) | -0.19(**) | 0.23(**) | -0.42(**) |
| Absb @ 400 | -0.23(**) | -0.01 | 0.10(*) | -0.06 | 0.05 | 0.21(**) | 0.35(**) | -0.19(**) | 0.23(**) | -0.42(**) |
| HIX | 0.06 | 0.02 | 0 | 0.11(*) | 0.13(**) | -0.07 | -0.14(**) | 0.24(**) | 0.01 | 0.10(*) |
| amino acids | -0.04 | -0.03 | 0.09 | -0.04 | 0.28(**) | 0.03 | 0.11(*) | -0.02 | 0.48(**) | -0.15(**) |
| TOC_L | -0.20(**) | 0.01 | 0.12(*) | -0.02 | 0.06 | 0.29(**) | 0.35(**) | -0.18(**) | 0.32(**) | -0.40(**) |
| TON_L | 0.18(**) | 0.02 | -0.08 | -0.05 | 0.24(**) | -0.14(**) | -0.21(**) | 0.78(**) | 0.11 (*) | 0.09 |
| CN_L | -0.25 (**) | -0.04 | 0.16(**) | -0.02 | 0.03 | 0.33(**) | 0.50(**) | -0.33(**) | -0.04 | -0.47(**) |
| BD | 0.46(**) | 0.05 | -0.42(**) | -0.22(**) | -0.07 | -0.51(**) | -0.83(**) | 0.35(**) | -0.14(**) | 0.70(**) |

**Table 2 continued**
| Variable | Ec | Phenols | Absob @<br>254 | Absob @<br>400 | HIX | amino<br>acids | TOC_L | TON_L | CN_L | BD |
| --- | --- | --- | --- | --- | --- | --- | --- | --- | --- | --- |
| **Ec** | 1 |  |  |  |  |  |  |  |  |  |
| **Phenols** | 0.04 | 1 |  |  |  |  |  |  |  |  |
| **Absb @ 254** | -0.04 | 0.58(**) | 1 |  |  |  |  |  |  |  |
| **Absb @ 400** | -0.08 | 0.60(**) | 0.97(**) | 1 |  |  |  |  |  |  |
| **HIX** | 0.37(**) | -0.09 | -0.04 | -0.22(**) | 1 |  |  |  |  |  |
| **amino acids** | -0.01 | 0.23(**) | 0.09 | 0.09 | 0.11(*) | 1 |  |  |  |  |
| **TOC_L** | 0.01 | 0.56(**) | 0.78(**) | 0.71(**) | 0.08 | 0.23(**) | 1 |  |  |  |
| **TON_L** | 0.66 (**) | -0.08 | -0.13(**) | -0.14(**) | 0.31 (**) | 0.05 | -0.05 | 1 |  |  |
| **CN_L** | -0.05 | 0.34(**) | 0.38(**) | 0.37(**) | -0.06 | 0.02 | 0.38(**) | -0.25 (**) | 1 |  |
| **BD** | 0.10(*) | -0.38(**) | -0.35 (**) | -0.33 (**) | 0.04 | -0.17 (**) | -0.37 (**) | 0.21(**) | -0.4 (**) | 1 |

**Note:** \*Correlation is significant at $P < 0.05$ level, and \*\* at the $P < 0.01$ level; $q_{\text{Mic}}$ , microbial quotient; $q_{\text{CO}_2}$ , metabolic quotient; Mic C, microbial carbon ( $\text{mg C kg}^{-1}$ ); Mic N, microbial nitrogen ( $\text{mg C kg}^{-1}$ ); Mic C:N, microbial C:N ratio; SR, soil respiration ( $\text{mg C kg}^{-1} \text{ h}^{-1}$ ); SOC, soil organic carbon ( $\text{mg C kg}^{-1}$ ); $\text{NO}_3^-$ , nitrate ( $\text{mg N l}^{-1}$ ); $\text{NH}_4^+$ , ammonium ( $\text{mg N l}^{-1}$ ); EC, ( $\mu\text{S cm}^{-1}$ ); Phenols, Soluble phenolics ( $\text{mg l}^{-1}$ ); Abs @ 254 and 400, absorbance of soil solution at 254 and 400 nm; HIX, humification index; Am acids, Free amino acids ( $\mu\text{M}$ ); TOC/N L, total organic carbon/nitrogen in leachate ( $\text{mg l}^{-1}$ ); BD, bulk density.

Due to differences in the units of individual variables, Factor Analysis (FA) was performed using a correlation matrix on the standardised values of the measured 20 attributes. The generalised least-squares method was used to extract factors because it is robust and requires no assumptions of sample coming from a multivariate normal distribution (SPSS, 2006). The first six factors with eigenvalues > 1 were retained for interpretation, whilst factors with eigenvalues < 1 explained less total variation than individual soil attributes (Brejda et al., 2000). The retained factors accounted for > 61% of the total variance in the measured attributes; see Table 3.

**Table 3.**
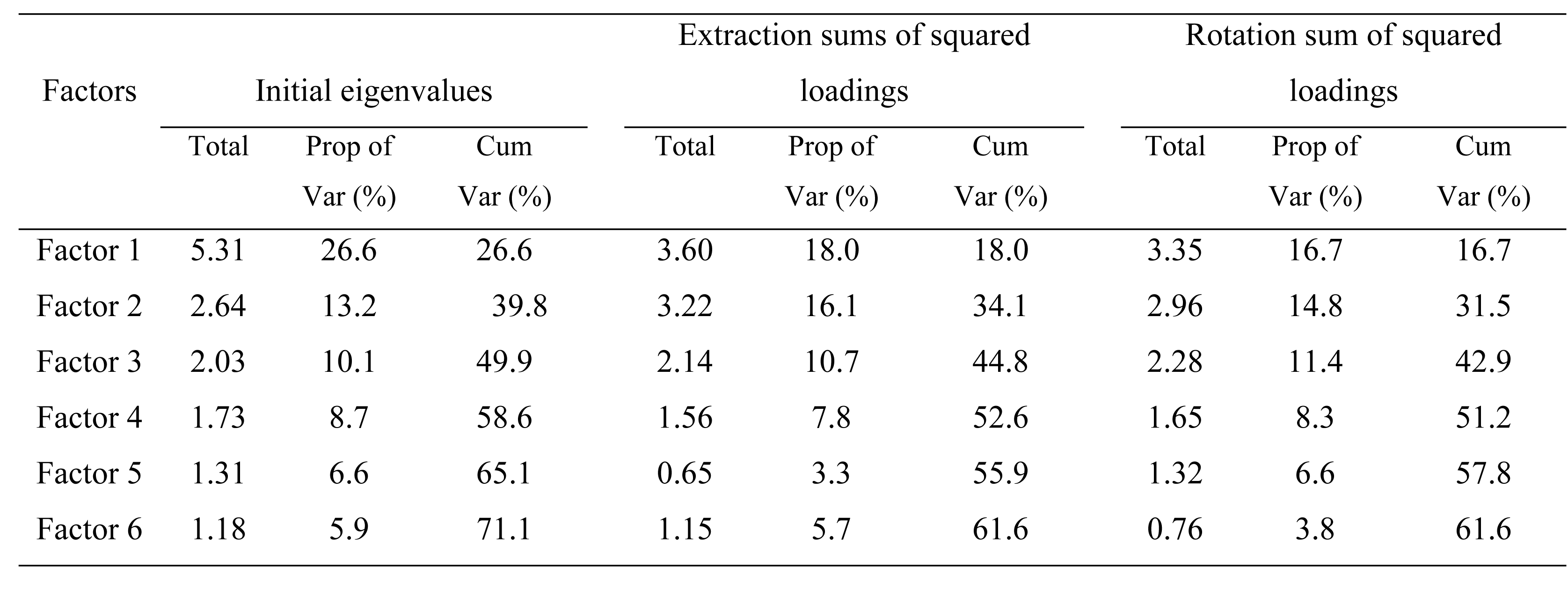
Total variance (Eigenvalue), proportion and cumulative variance (Prop Var and Cum Var) explained by factor analysis using correlation matrix (standardized data) on the measured attributes.

| Factors | Initial eigenvalues |  |  | Extraction sums of squared |  |  | Rotation sum of squared |  |  |
| --- | --- | --- | --- | --- | --- | --- | --- | --- | --- |
|  |  |  |  | loadings |  |  | loadings |  |  |
|  | Total | Prop of<br>Var (%) | Cum<br>Var (%) | Total | Prop of<br>Var (%) | Cum<br>Var (%) | Total | Prop of<br>Var (%) | Cum<br>Var (%) |
| Factor 1 | 5.31 | 26.6 | 26.6 | 3.60 | 18.0 | 18.0 | 3.35 | 16.7 | 16.7 |
| Factor 2 | 2.64 | 13.2 | 39.8 | 3.22 | 16.1 | 34.1 | 2.96 | 14.8 | 31.5 |
| Factor 3 | 2.03 | 10.1 | 49.9 | 2.14 | 10.7 | 44.8 | 2.28 | 11.4 | 42.9 |
| Factor 4 | 1.73 | 8.7 | 58.6 | 1.56 | 7.8 | 52.6 | 1.65 | 8.3 | 51.2 |
| Factor 5 | 1.31 | 6.6 | 65.1 | 0.65 | 3.3 | 55.9 | 1.32 | 6.6 | 57.8 |
| Factor 6 | 1.18 | 5.9 | 71.1 | 1.15 | 5.7 | 61.6 | 0.76 | 3.8 | 61.6 |

The retained factors were subjected to a varimax rotation. A varimax rotation redistributes the variance of significant factors and minimizes the number of variables that have high loadings on each factor, thereby simplifying the interpretation of the factors (SPSS, 2006). The relative importance of each soil attribute, in terms of its contribution to all of the factors, was judged by its communality value (Field, 2005; Ayoubi and Khormali, 2008) and is shown in Table 4. The six factors explained > 90% variance in absorbance @ 254 and 400 (absb@254 and 400), microbial carbon (Mic C), and soil organic carbon (SOC); > 80% in total organic nitrogen in leachate (TON_L) and bulk density (BD); > 70% in microbial nitrogen (Mic N), Nitrate, Ammonium, electrical conductivity (EC), and total organic carbon in leachate (TOC_L); > 60 % in microbial quotient (*q*Mic), pH and humification index (HIX); > 50 % microbial C/N ratio (Mic CN), soil respiration (SR), and phenolics; and < 50 % C/N ratio of the leachate (CN_L) and microbial metabolic quotient (*q*CO_2_) (Table 4). Attributes with the low communality estimates (e.g. *q*CO_2_ and leachate C:N) were the least important for interpreting factors. The magnitudes of the loadings were used as a criterion for interpreting the relationship between the soil attributes and the factors. Soil attributes were assigned to the factor for which the loadings were highest.

**Table 4.** Proportion of variance (loadings) using varimax rotation and communality estimates for soil attributes of the retained factors.

| Variable | Factor 1 | Factor 2 | Factor 3 | Factor 4 | Factor 5 | Factor 6 | Communality extraction |
| --- | --- | --- | --- | --- | --- | --- | --- |
| *q*Mic | -0.54 | -0.13 | 0.12 | 0.45 | -0.01 | -0.07 | 0.67 |
| *q*CO<sub>2</sub> | 0.05 | -0.05 | -0.05 | -0.18 | 0.01 | 0.00 | 0.10 |
| Microbial-C | 0.29 | 0.03 | -0.04 | 0.89 | 0.09 | -0.03 | 0.90 |
| Microbial-N | 0.05 | -0.04 | -0.08 | 0.75 | -0.02 | 0.14 | 0.73 |
| Microbial C:N | 0.07 | 0.05 | 0.30 | -0.03 | 0.11 | 0.01 | 0.51 |
| Soil respiration | 0.61 | 0.06 | -0.07 | 0.09 | 0.03 | -0.06 | 0.50 |
| Soil organic C | 0.92 | 0.16 | -0.08 | 0.08 | 0.01 | -0.06 | 0.91 |
| Nitrate-N | -0.27 | -0.09 | 0.81 | -0.04 | 0.01 | 0.04 | 0.77 |
| Ammonium-N | 0.01 | 0.23 | 0.05 | 0.05 | 0.78 | 0.01 | 0.72 |
| pH | -0.68 | -0.28 | 0.02 | -0.06 | -0.18 | 0.05 | 0.68 |
| Elec. conductivity | 0.03 | 0.00 | 0.74 | -0.03 | -0.05 | 0.22 | 0.70 |
| Soluble phenolics | 0.29 | 0.52 | -0.03 | 0.06 | 0.32 | -0.10 | 0.55 |
| Absorb @ 254 nm | 0.17 | 0.98 | -0.06 | -0.01 | 0.04 | 0.03 | 1.00 |
| Absorb @ 400 nm | 0.17 | 0.96 | -0.05 | -0.02 | 0.04 | -0.20 | 0.99 |
| HIX | -0.06 | -0.06 | 0.25 | 0.07 | 0.05 | 0.76 | 0.69 |
| Amino acids | 0.11 | 0.06 | 0.02 | 0.01 | 0.66 | 0.05 | 0.56 |
| TOC (leachate) | 0.24 | 0.71 | -0.02 | 0.00 | 0.29 | 0.18 | 0.73 |
| TON (leachate) | -0.12 | -0.06 | 0.91 | 0.01 | 0.09 | 0.07 | 0.87 |
| C:N (leachate) | 0.47 | 0.26 | -0.17 | -0.02 | -0.06 | 0.02 | 0.42 |
| Bulk density | -0.86 | -0.19 | 0.13 | -0.18 | -0.11 | -0.05 | 0.87 |

The **first factor** explained 16.7 % (see Table 3) of the total variance. It was named *soil organic matter (SOM)* because it had high positive loading for SOC (0.92), soil respiration (0.61) and leachate C:N ratio (47), a high negative loadings for bulk density (−0.86), pH (−0.68) and moderately on *q*Mic (−0.54). Grouping *q*Mic with the *SOM* factor rather than factor 4 was as a result of its stronger correlation with attributes constituting the *SOM* factor namely, soil respiration (*r* = −0.26), SOC (*r* = −0.47) and bulk density (*r* = 0.46) rather than with Microbial-C (*r* = 0.23) and Microbial-N (*r* = 0.17) of factor 4 (Table 3). **The second factor** explained 15% of the total variance with a high positive loading for soluble phenolics (0.52), leachate absorbance at 254 nm (0.98), 400 nm (0.96) and leachate TOC (0.71) and consequently, was termed *OM humification.* The **third factor** explained 11 % of the total variance with high positive loadings for nitrate (0.81), leachate TON (0.91) and electrical conductivity (0.74) and was therefore termed *soluble nitrogen* factor. The **fourth factor** had positive loadings for Microbial-C (0.89), Microbial-N (0.75) and a moderately high loading for *q*Mic (0.45), and was termed *microbial biomass*. The **fifth factor** had positive loading for ammonium (0.78) and amino acids (0.66) and was termed *reduced N*. **The sixth factor** explained only 4 % of the total variance and had a high positive loading for HIX (0.76) and was termed *soil humification index*.

### 3.3. Effect of soil types on attribute means and factor scores

One way ANOVA revealed that most of the soil attributes and factor scores varied significantly with soil types (Table 5). However, pairwise comparison showed that the effect of soil types on most attribute was very small. In most cases, only the Peat soils were clearly significantly (*P <* 0.01) different from all the other soil types. Only *SOM* and *microbial biomass* factors (Factors 1 and 4 respectively) varied significantly (*P <* 0.05) with soil type. *SOM* factor mean scores were negative for Brown, GWG, SWG and Pelosol soils and positive for Lithomorphic, Peat and Podzolic soils. Peats had the highest score and were significantly different from all other soil types on the *SOM* factor. Furthermore, Peat soils had the highest SOC content to which the analysis also confirmed. The mean scores for *SOM* factor did not vary significantly (*p* > 0.05) within Browns, GWGs and Pelosols nor did it do so among the Lithomorphic, Podzolic and SWG soils. The *Microbial biomass* factor varied significantly (*P <* 0.05) between Brown versus GWG soil types and Lithomorphics only. Mean scores for *OM humification*, *soluble N*, *reduced N* and *humification index* did not vary significantly (*P* > 0.05) among all soil types.

**Table 5.** Soil attribute means and factor scores in the different soil types (The first 5 variables are the most important for discrimination between soil types)

| Soil attributes | Soil types |  |  |  |  |  |  | SEM | ANOVA |
| --- | --- | --- | --- | --- | --- | --- | --- | --- | --- |
|  | Brown | Groundwater<br>gley | Lithomorphic | Peat | Pelosols | Podzolic | Surface<br>water gley |  |  |
| Microbial quotient | 0.018<sup>a</sup> | 0.026<sup>a</sup> | 0.014<sup>ac</sup> | 0.003<sup>b</sup> | 0.014<sup>abc</sup> | 0.010<sup>c</sup> | 0.018<sup>a</sup> | 0.003 | 0.00 |
| *q*CO<sub>2</sub> | 0.073 | 0.002 | 0.001 | 0.011 | 0.01 | 0.002 | 0.003 | 0.012 | NS |
| Microbial-C (g kg<sup>-1</sup>) | 0.59<sup>a</sup> | 1.00<sup>ab</sup> | 1.03<sup>ab</sup> | 1.37<sup>b</sup> | 0.54<sup>a</sup> | 1.02<sup>ab</sup> | 0.89<sup>ab</sup> | 0.13 | 0.00 |
| Microbial-N (mg kg<sup>-1</sup>) | 85<sup>a</sup> | 119<sup>ab</sup> | 148<sup>b</sup> | 113<sup>ab</sup> | 71<sup>ab</sup> | 111<sup>ab</sup> | 99<sup>ab</sup> | 16 | 0.03 |
| C:N (Microbial) | 12.4 | 19.6 | 18.9 | 19.7 | 36.3 | 29.9 | 33.2 | 12 | NS |
| Soil respiration (mg kg<sup>-1</sup> h<sup>-1</sup>) | 0.63<sup>a</sup> | 1.10<sup>a</sup> | 0.93<sup>a</sup> | 3.35<sup>b</sup> | 1.63<sup>ab</sup> | 1.58<sup>ab</sup> | 1.18<sup>a</sup> | 0.45 | 0.00 |
| Soil organic C (g kg<sup>-1</sup>) | 42<sup>a</sup> | 45<sup>a</sup> | 132<sup>b</sup> | 377<sup>c</sup> | 92<sup>ab</sup> | 151<sup>b</sup> | 98<sup>b</sup> | 23 | 0.00 |
| Nitrate (mg N l<sup>-1</sup>) | 3.00<sup>a</sup> | 2.04<sup>ac</sup> | 2.32<sup>ac</sup> | 0.13<sup>b</sup> | 1.13<sup>c</sup> | 0.37<sup>bc</sup> | 3.08<sup>a</sup> | 0.39 | 0.00 |
| Ammonium (mg N l<sup>-1</sup>) | 0.25 | 0.18 | 0.3 | 0.27 | 0.17 | 0.31 | 0.3 | 0.05 | NS |
| pH | 6.55<sup>a</sup> | 6.56<sup>a</sup> | 6.24<sup>ac</sup> | 4.71<sup>b</sup> | 6.18<sup>ac</sup> | 5.08<sup>b</sup> | 5.73<sup>c</sup> | 0.2 | 0.00 |
| Elect. conductivity (µS cm<sup>-1</sup>) | 129 | 107 | 124 | 99 | 74 | 81 | 116 | 16 | NS |
| Soluble phenols (mg l<sup>-1</sup>) | 0.33<sup>ac</sup> | 0.26<sup>a</sup> | 0.68<sup>bc</sup> | 1.10<sup>b</sup> | 0.56<sup>abc</sup> | 1.20<sup>b</sup> | 0.46<sup>c</sup> | 0.16 | 0.00 |
| Absorbance @ 254 nm | 0.25<sup>a</sup> | 0.28<sup>a</sup> | 0.29<sup>ab</sup> | 0.47<sup>b</sup> | 0.45<sup>ab</sup> | 0.48<sup>b</sup> | 0.32<sup>ab</sup> | 0.48 | 0.00 |
| Absorbance @ 400 nm | 0.028<sup>a</sup> | 0.033<sup>a</sup> | 0.032<sup>ab</sup> | 0.061<sup>b</sup> | 0.047<sup>ab</sup> | 0.061<sup>b</sup> | 0.036<sup>ab</sup> | 0.009 | 0.00 |
| Humification index (HIX) | 9.0<sup>ab</sup> | 9.0<sup>ab</sup> | 8.7<sup>ab</sup> | 8.2<sup>a</sup> | 8.3<sup>ab</sup> | 8.6<sup>ab</sup> | 9.3<sup>b</sup> | 0.3 | 0.03 |
| Amino acids (µM) | 1.52 | 1.83 | 1.67 | 1.95 | 1.15 | 3.1 | 2.08 | 0.4 | NS |
| Leachate TOC (mg l<sup>-1</sup>) | 7.5<sup>a</sup> | 6.9<sup>a</sup> | 8.2<sup>ab</sup> | 12.0<sup>b</sup> | 12.8<sup>ab</sup> | 12.3<sup>b</sup> | 9.8<sup>ab</sup> | 2.2 | 0.00 |
| Leachate TON (mg l<sup>-1</sup>) | 5.82<sup>a</sup> | 3.47<sup>ac</sup> | 3.16<sup>ac</sup> | 0.78<sup>b</sup> | 1.62<sup>c</sup> | 1.81<sup>bc</sup> | 6.69<sup>a</sup> | 0.8 | 0.01 |
| Leachate C:N | 4.6<sup>a</sup> | 5.5<sup>a</sup> | 7.2<sup>a</sup> | 19.0<sup>b</sup> | 9.1<sup>ab</sup> | 17.5<sup>b</sup> | 9.7<sup>a</sup> | 2.4 | 0.00 |
| Bulk density | 1.10<sup>a</sup> | 1.11<sup>a</sup> | 0.63<sup>b</sup> | 0.19<sup>c</sup> | 1.08<sup>a</sup> | 0.58<sup>b</sup> | 0.81<sup>b</sup> | 0.06 | 0.00 |
| **Factors** | **Factor scores** |  |  |  |  |  |  |  |  |
| Factor 1 | -0.52<sup>a</sup> | -0.63<sup>a</sup> | 0.15<sup>b</sup> | 1.58<sup>c</sup> | -0.59<sup>a</sup> | 0.2<sup>b</sup> | -0.07<sup>b</sup> | 0.12 | 0 |
| Factor 2 | -0.17 | -0.05 | -0.13 | 0.22 | -0.46 | 0.44 | 0 | 0.15 | NS |
| Factor 3 | 0.09 | -0.1 | -0.06 | -0.13 | -0.4 | -0.28 | 0.23 | 0.11 | NS |
| Factor 4 | -0.24<sup>a</sup> | 0.36<sup>b</sup> | 0.30<sup>b</sup> | 0.03<sup>ab</sup> | -0.21<sup>ab</sup> | 0.01<sup>ab</sup> | 0.02<sup>ab</sup> | 0.19 | 0.04 |
| Factor 5 | -0.05 | -0.22 | -0.03 | -0.11 | -0.2 | 0.36 | 0.14 | 0.17 | NS |
| Factor 6 | 0.06 | -0.1 | 0.17 | -0.3 | -0.29 | -0.2 | 0.23 | 0.18 | NS |

### 3.4. Soil quality indicators across soil types

Discriminant analysis of the six statistical factors in relation to soil types, indicated that the *SOM* was the most powerful in discriminating among the seven soil type groups based on the magnitude of their discriminant coefficients (Eq. 2). The first canonical discriminant function explained 90 % of the total variance based on Wilks’s Lambda, (*P <* 0.001) (table not shown) and therefore was the most important canonical discriminant function for discriminating soil types using the soil quality factors identified. Although the second canonical discriminant function was also significant (*P* = 0.03) based on Wilks’s Lambda, it only accounted for 4 % of the total variance and therefore was not used.

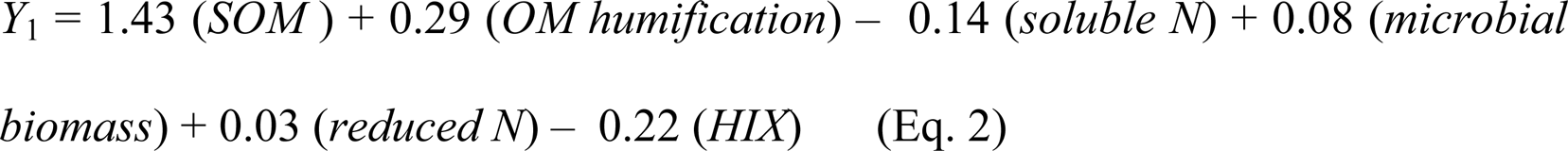

Therefore the group differences across soil types shown by ANOVA can be explained in terms of *SOM*, judging from the discriminant coefficient which was five-fold larger than the coefficient for the *OM humification* factor and several fold greater than the rest of the factors. Discriminant analysis of the measured attributes constituting *SOM* (i.e. *q*Mic, soil respiration (SR), soil organic C (SOC), pH and bulk density (BD)) indicated that microbial quotient (*q*Mic) was the most powerful attribute discriminating the soil types (Eq. 3).

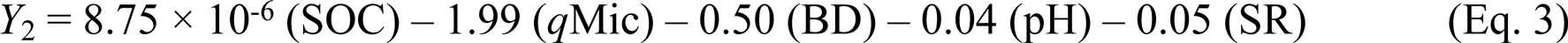

The discriminant coefficient for *q*Mic was four-fold larger than the coefficient for bulk density and more than 40-fold for the rest. The *q*Mic was significantly correlated with bulk density (0.46**), soil organic C (−0.47**), pH (0.35**) and soil respiration (−0.26**) while bulk density was significantly correlated with soil organic C (−0.83**), pH (0.70**) and soil respiration (−0.53**) meaning that *q*Mic and bulk density, though correlated, were the most important and dominant attributes for assessing soil quality across soil types. The mean comparisons using the Games-Howell approach indicated that the bulk density had similar discriminating power as the *SOM* factor among the soil types. *q*Mic mean values varied significantly with soil types separating Peat < Podzols < Browns, GWGs and SWGs soils in increasing order (Table 5).

### 3.5. Effect of aggregate vegetation class on factor scores

Aggregate vegetation class (AVC) showed more effects on factor scores than the soil types. The significant effects were observed in *SOM*, *OM humification*, *microbial biomass* and *humification index*. The *soluble N* and *reduced N* factors showed no significant variation among the AVCs (Table 6). The *SOM* factor had the highest factor scores (*P <* 0.001) in Heath and Bog. Mean scores between Moorland Grass Mosaics and Upland Woodland did not vary significantly (*P* > 0.05); nor among Fertile Grasslands, Infertile Grassland, Lowland Woodland and Tall Grass Mosaic. The mean scores were lowest in Crop and Weeds and were significantly different (*P <* 0.001) from all other AVCs except in Tall Grass and Herbs.

**Table 6.** Effect of Aggregate Vegetation Class (AVC) on factor scores and soil attribute means.

| Average vegetation class mean factor scores |  |  |  |  |  |  |  |  |  |  |
| --- | --- | --- | --- | --- | --- | --- | --- | --- | --- | --- |
| Factors | Crops &<br>weeds | Fertile<br>grasslands | Heath<br>& bog | Infertile<br>grassland | Lowland<br>woodland | Moorland grass<br>mosaics | Tall grass<br>& herbs | Upland<br>woodland | SEM | ANOVA |
| **Factor 1** | -0.80<sup>a</sup> | -0.54<sup>b</sup> | 1.43<sup>c</sup> | -0.50<sup>b</sup> | -0.40<sup>b</sup> | 0.62<sup>d</sup> | -0.64<sup>ab</sup> | 0.20<sup>bd</sup> | 0.10 | 0.00 |
| **Factor 2** | -0.40<sup>a</sup> | -0.11<sup>ab</sup> | 0.30<sup>b</sup> | 0.02<sup>b</sup> | 0.41<sup>ab</sup> | -0.11<sup>ab</sup> | -0.06<sup>ab</sup> | 0.51<sup>ab</sup> | 0.19 | 0.00 |
| **Factor 3** | 0.34 | 0.07 | -0.09 | -0.03 | 0.05 | -0.34 | 0.12 | -0.28 | 0.14 | NS |
| **Factor 4** | -0.49<sup>a</sup> | 0.16<sup>b</sup> | 0.07<sup>b</sup> | 0.27<sup>b</sup> | -0.19<sup>ab</sup> | 0.28<sup>b</sup> | -0.61<sup>a</sup> | -0.21<sup>ab</sup> | 0.16 | 0.00 |
| **Factor 5** | -0.39 | 0.09 | 0.13 | 0.02 | -0.12 | 0.22 | -0.20 | 0.18 | 0.14 | NS |
| **Factor 6** | -0.29<sup>a</sup> | 0.04<sup>ab</sup> | -0.35<sup>ab</sup> | 0.13<sup>b</sup> | 1.15<sup>c</sup> | 0.18<sup>b</sup> | 0.38<sup>bc</sup> | 0.63<sup>bc</sup> | 0.17 | 0.00 |
| **<u>Soil</u>** |  |  |  |  |  |  |  |  |  |  |
| **<u>attributes</u>** |  |  |  |  |  |  |  |  |  |  |
| **<u>Soil attribute mean values</u>** |  |  |  |  |  |  |  |  |  |  |
| **Soil respiration** | 0.29<sup>a</sup> | 1.00<sup>b</sup> | 3.22<sup>c</sup> | 0.77<sup>b</sup> | 0.67<sup>ab</sup> | 1.44<sup>b</sup> | 0.43<sup>ab</sup> | 1.41<sup>b</sup> | 0.23 | 0.000 |
| **Soil organic C** | 16.7<sup>a</sup> | 43.6<sup>b</sup> | 350.2<sup>c</sup> | 43.8<sup>b</sup> | 46.4<sup>b</sup> | 185.6<sup>c</sup> | 25.0<sup>ab</sup> | 119.8<sup>c</sup> | 11.2 | 0.000 |
| **pH** | 7.3<sup>a</sup> | 6.4<sup>b</sup> | 4.6<sup>c</sup> | 6.3<sup>b</sup> | 6.2<sup>abd</sup> | 5.2<sup>d</sup> | 6.6<sup>ab</sup> | 4.7<sup>dc</sup> | 0.2 | 0.000 |
| **Bulk density** | 1.37<sup>a</sup> | 1.06<sup>b</sup> | 0.21<sup>c</sup> | 0.95<sup>b</sup> | 0.89<sup>b</sup> | 0.41<sup>d</sup> | 1.22<sup>ab</sup> | 0.48<sup>d</sup> | 0.05 | 0.000 |
| **qMic** | 0.021<sup>a</sup> | 0.023<sup>a</sup> | 0.005<sup>b</sup> | 0.021<sup>a</sup> | 0.015<sup>ab</sup> | 0.009<sup>b</sup> | 0.015<sup>ab</sup> | 0.010<sup>ab</sup> | 0.003 | 0.000 |

Means scores for *OM humification* factor varied significantly (*P <* 0.001) between Crop and Weeds verses Herb and Bog, and Infertile Grasslands; all other pairs did not vary significantly. For *microbial biomass* factor, Crop and Weeds and Tall Grass and Herbs varied significantly (*P <* 0.001) against the Fertile Grassland, Infertile Grasslands, Heath and Bog, and Moorland Grass Mosaics, while all other pairs were not significantly different (*P* > 0.05). The *humification index* factor showed that the mean scores varied significantly (*P <* 0.001) among Crop and Weeds versus Infertile Grassland and Moorland Grass Mosaics versus Lowland Woodland only.

### 3.6. Soil quality indicators across Aggregate Vegetation Classes (AVC)

The first canonical discriminant function of the discriminant analysis of the six factors across the AVCs explained 94% of the total variance (Wilks’s Lambda, *P <* 0.001) whose coefficients were used in the equation below:

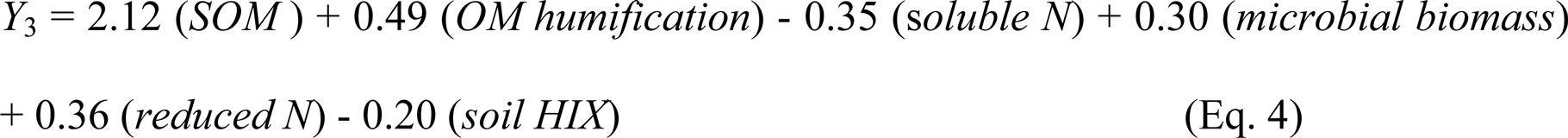

From the discriminant coefficients in Eq. [4], *SOM* factor was the most powerful discriminating among the eight different AVCs. The *SOM* factor was more than four-fold larger than the coefficients of all others soil quality factors under consideration.

The discriminant analysis of the measured attributes constituting the *SOM* factor showed that BD and *q*Mic were the most powerful discriminating soil attributes among the seven habitats (AVCs) (Eq. 5).

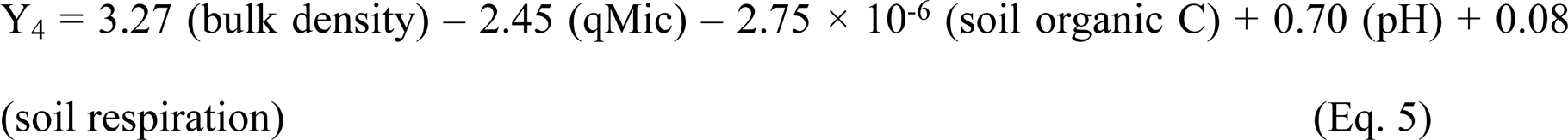

Bulk density possessed similar discriminating power as the *SOM* factor among the AVCs. Bulk density values were significantly different (*P <* 0.001) among AVCs with the lowest mean values in Heath and Bog (0.21 g cm^-3^) < Upland Wooded (0.48 g cm^-3^) and Moorland Grass Mosaic (0.41 g cm^-3^) < Fertile Grass (1.06 g cm^-3^), Infertile Grass (0.95 g cm^-3^), Lowland Wooded (0.89 g cm^-3^) < Tall Grass and Herbs (1.21 g cm^-3^) and Crop and Weeds (1.37 g cm^-3^; Table 6).

### 3.7. Main and interactions effect of soil types and AVCs

The results of the two-way ANOVA on the first canonical discriminate function on all 20 variables showed significant (*P <* 0.01) main and interaction effects. The main effect of soil types and the effect of soil types * AVCs interaction on the attribute’s scores was very small (Partial Eta Square = 0.09 and 0.16 respectively), while the main effect of the AVCs was large (Partial Eta Square = 0.42; Table. 7).

The cross tabulation of AVCs versus soil types (Table S3), showed that 27 out of 56 combinations or cells, the soil types were sampled less than the calculated expected counts in the AVCs. In 16 combinations, the soil types were not at all represented in the AVCs. The most affected were the Lowland Woodland and Tall Grass and Herbs where, only Brown and SWGs were samples in the former and only Browns, GWG and SWGs in the latter.

## 4. Discussion

### 4.1. Effect of soil types and AVCs on the soil quality factors and/or indicators

A set of 20 correlated soil attributes were grouped into six factors called soil quality factors, using factor analysis. The factors identified contribute to one or more key soil functions proposed by Larson and Pierce (1991) and therefore could be considered soil quality indicators (Brejda et al., 2000). Since the soil quality factors cannot be measured directly (Elliott, 1997; Brejda et al., 2000), the effect of soil types and the AVCs on these factors were inferred by monitoring soil attributes that comprised them.

Not all the soil quality factors varied significantly with soil types or with AVCs. Only *SOM* and *microbial biomass* factors varied significantly (*P <* 0.001) by soil types. *SOM* was able to discriminate the highest number of soil groups, separating the Peats (1) with the highest scores, from Lithomorphics, Podzols, and SWGs (2) with intermediate scores, and from Browns, GWGs and Pelosols (3) with the lowest scores (Fig. 5), thus rendering three distinct soil type groupings. The *microbial biomass* factor had a minor effect, discriminating the Browns from GWGs and Lithomorphics only. The soil attributes constituting these soil quality factors (Soil respiration, SOC, pH, bulk density, *q*Mic, Microbial-C and -N) showed significant (*P* < 0.01) variations discriminating at most three groups of soil types. In all the attributes considered, Browns, GWGs and Pelosols were grouped together. *SOM* factor, SOC and bulk density attributes separated the Peats as a unique soil group from all other soil types, which is not entirely a surprising result, since the peats are highly organic in nature with low BD as opposed to mineral soils with low OM content and higher bulk densities. The most important soil quality indicator associated with specific soil types or groups was the *SOM* factor with *q*Mic > bulk density as the most important attributes.

Similarly, the most important SQF differentiating the AVCs across the GB was *SOM* factor with bulk density > *q*Mic attributes being the most important attributes. Four distinct AVC groups were separated based on *SOM* factor and BD. Heath and bog was exclusively separated as one group (1). Other groups were: (2) Crop and weeds with Tall grass and herb; (3) Fertile grassland, Infertile grassland, Lowland woodland, Tall grass and herbs, and Upland woodland; (4) Moorland grassland mosaic with Upland woodland. The Upland woodland and Tall grass and herbs were intermediate habitats classifying in more than one of these groups. The rest of the factors and attributes discriminated three or less groups. The soil attributes were generally better in discriminating the AVCs than the SQF (Table 6)

Since *q*Mic and bulk density were moderately correlated (*r=*0.46**), they may be redundant as indicators to be used together. If only one attribute were to be used to monitor soil quality in soil types and AVCs, *q*Mic and BD respectively seems to offers the greatest potential judging from their high weights on the respective prediction models. However the *q*Mic may be a ‘MUST be included’ soil attribute in the minimum data set, due to its important role in several soil functions, being a fraction of soil carbon. Soil C influences a wide range of soil functions including bulk density, infiltration, pesticide buffering, aeration, aggregate formation, pH, buffer capacity, cation-exchange properties, mineralization, and the activity of soil organisms (Larson and Pierce, 1991; Seybold et al., 1997). However, since the measurement of bulk density is reasonably easy to obtain, it is therefore reasonable to consider it together with SOC, microbial and biomass C as minimum data set for assessing soil quality across average vegetation classes in the study area.

Pedogenesis has taken place over thousands of years in the UK. During this period there has been a range of climate change related vegetation colonization phases starting from tundra heath and cycling through a range of forest types (Fitzpatrick, 1980). During this period parent material/topography, climate and vegetation would have been stable for long periods of time leading to the differentiation of soils. This was followed by progressive forest clearance which started approximately 1000-3000 years ago with vegetation cover becoming more grassland and heathland dominated. The last 200 years, however, has seen intense management of these soils with the addition of fertilisers, lime and organic wastes combined with mechanical mixing of the soils which has reversed centuries of acidification and soil horizon development. This homogenisation of the soil has led to shifts between soil types even over short timescales (e.g. humic-podzolic to brown soils on improved upland grasslands) and the loss of peat soils in intensive agricultural areas (e.g. East Anglia; Taft et al., 2018). One key question is therefore whether it is historical soil type or current vegetation that is more important in driving soil processes in the short term (e.g. over a 10-25 year timescale)? Here we found that more soil quality factors showed an AVC effect rather than a soil type effect. All soil quality factors varied significantly (*P* < 0.01) by AVCs except *soluble N* and *reduced N* factors though none discriminated more than four groups. It is possible that some of the soil quality factors that were insensitive to vegetation may represent inherent soil qualities that are controlled by other key factors of soil formation (e.g. parent material/topography), while those which significantly varied by AVCs may represent dynamic soil qualities, possessing great potential for assessing management practices on soil quality (Jenny, 1994; Soybold et al., 1997; Brejda et al., 2000; Bünemann, et al., 2018).

Most indicators available in literature have not been validated nor their sensitivity tested in a wide range of situations (Velasquez et al., 2007). Some of the attributes measured and the soil quality indicators identified in this work are not usually used in the monitoring of soil quality, but are candidates for potential alternatives (Schloter, et al., 2018).

### 4.2. Prediction of SQF and SQI by soil type or AVC

The clusters from multivariate classification are “natural” groups, which uses the “minimum-variance” solution; where a population is partitioned into cluster subsets by minimizing the total within group variation while maximising between groups variance (Wishart, 1968). The groups/cluster formed from the multivariate analysis need to have no significant overall spread. The clusters therefore, should correspond to data modes (distinct modes). However, most of our cluster modes defined by soil types were not always distinct. Most of them were separated from each other by significant “noise” data, making it impossible to resolve all the clusters. Thus, the definition of the reference values for each soil type or AVC was ambiguous, since most soils types or AVC groups could not be differentiated (Fig. 2). Forming, describing and defining the groups could involve the use all measured attributes even though only a few could be differentiating (Soil Survey Staff, 1999). Even when the soil quality factors/indicators and attributes were used in combination, some groups/clusters could still not be resolved. Therefore, the soil quality indicators and attributes identified in this study can only be used to characterise soil types and AVCs groups rather than for prediction or classification. From the discriminant plots and the dendrogram in Fig. 3 three groups can be defined in soil types and four groups in the AVC.

**Figure 2.**
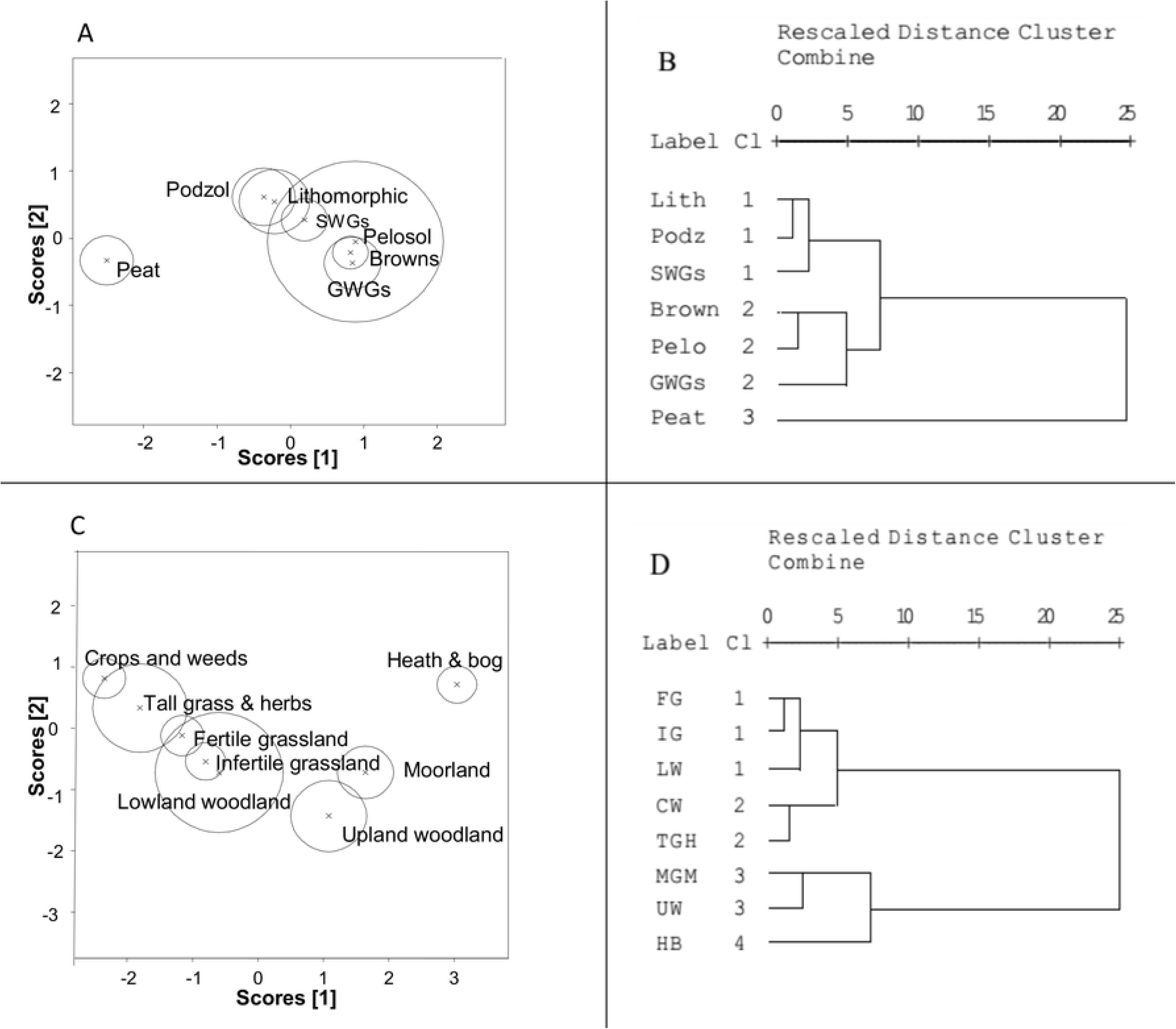
Discrimination plots showing 95% confidence circles around the means for soil types (Panel A) and AVCs (Panel C). Panels B and D are the respective cluster analyses dendrograms using a complete linkage method.

Defining differentiating criteria for these groups in the soil types could involve the use of bulk density attribute to define unique property ranges for the first groups, a combination of soil respiration and SOC attributes for the second group, and a combination of *q*Mic, soil respiration, SOC, pH and bulk density attributes for the third group. The Pelosols were the most dispersed and unreliable group for the purpose of attribute membership prediction, probably due to the fact that they were under sampled, considering that only six samples were included in the analysis. The classification of the AVCs using discriminate and cluster analyses on key attributes yielded four clusters. Defining differentiating criteria for these groups could involve the use of a combination of soil respiration, SOC, pH attributes to define property ranges for the first, second and fourth groups and bulk density attribute for the third group. Tall Grass and Herbs and Lowland Wooded were under sampled (with 11 and 6 samples respectively; Table S3) which greatly compromised their predictive accuracy as can be observed from the large 95% confidence circles which overlapped with other AVC groups.

### 4.3. To what extent do soil types and/ or AVC act as major regulators of SQI?

The two-way ANOVA and the tests of between-subjects effects on the first canonical discriminant (CD) function from the canonical discriminant analysis (CDA) of the 20 physical, chemical and biological properties showed significant differences between groups (soil types and AVCs) as well as significant differences in the effect of soil type on the soil attributes between the AVC (significant interaction of soil type × AVC; Table 7). The ‘practical’ significance of each term from Partial Eta Square values indicates that AVCs (with a large Partial Eta Square = 0.42), were a better regulator of the SQIs than soil types (with a weak Partial Eta Square = 0.09). The effect size for the interaction was equally relatively weak (Partial Eta Square = 0.16). The conclusion of the significant (*P <* 0.01) interaction effect of soil type × AVC is that the soil type differences in the first CD function (or attributes) partly depended on the AVCs where the soil was sampled. A multiple comparison of all soil type groups with AVC groups would be required to draw specific conclusions regarding the interaction effects, which is quite complex and is beyond the scope of this thesis. Suffice to say that there was a partial and varied soil type × AVC interaction across all levels. These interactions confirm Jenny’s (1994) theory that the biotic factor (of which vegetation plays a major role) is amongst others an important soil forming factor. However, the results from the cross tabulation indicated that not all soil types were well represented in the AVCs in going by the calculated expected counts. In some cases soil types were not at all represented (See Table S3). This problem can contribute to the complexity and accuracy in the interpretation of the interaction effect observed above.

**Table 7.** Tests of between-subjects effects;

| Source | Type IV sum of squares | Df | Mean Square | F | Sig. | Partial eta squared |
| --- | --- | --- | --- | --- | --- | --- |
| Corrected model | 553.14 | 38 | 14.56 | 36.36 | 0.001 | 0.844 |
| Intercept | 3.42 | 1 | 3.416 | 8.532 | 0.004 | 0.032 |
| **Soil Type \* AVC_Desc** | **18.98** | **25** | **0.759** | **1.896** | **0.008** | **0.157** |
| **Soil Type** | **10.36** | **6** | **1.726** | **4.311** | **0.001** | **0.092** |
| **AVC_Desc** | **73.97** | **7** | **10.57** | **26.39** | **0.001** | **0.420** |
| Error | 102.09 | 255 | 0.400 |  |  |  |
| Total | 655.24 | 294 |  |  |  |  |
| Corrected Total | 655.24 | 293 |  |  |  |  |

*Notes: Dependent variable: 1st Canonical Discriminant function scores from of all the soil attributes measured. AVC\_desc means AVC description; Soil Type\*AVC\_Desc means the interaction between the soil type and AVC effects.*

## 5. Conclusions

The dominant SQFs/Is and attributes varied by both soil type and AVC. The *SOM* factor was the most discriminating factor for both soil types and AVCs with microbial quotient and bulk density as the most discriminating measured attributes. The discriminant analysis on the important measured attributes comprising the *SOM* factor produced three fairly homogenous groups for soil types and four groups for AVCs. It was however, impossible to define reference values in the SQF/I or attributes for separate individual soil types or AVCs, as property ranges greatly overlapped due to large between group variability (probably due to integrating large spatial areas). Some of the differences observed in soil types with regard to soil attributes were in part dependent on the AVCs differences.

Therefore, whether SQIs can be predicted by soil types remains an open question. This study has shown that soil types or AVCs are poor predictors for SQF and indicators derived from factor analysis. However, different sets of SQIs and attributes for different regions have been used in the past in different studies (e.g. Brejda et al., 2000a; Brejda et al., 2000b; Shukla et al., 2006; Valesquez et al., 2007; Ayoubi and Khormali, 2008) suggesting that there may not be a universal optimum set of indicators for use across different regions of differing climatic conditions. Therefore, the search for SQIs which can be predicted by soil types continues.

For future work it might be worthwhile to make special consideration for the climatic, spatial and parent material variability in the sampling designs in addition to the inclusion of other promising soil attributes. Management factors should also be included (e.g. fertilizer regime). In terms of other key soil quality indicators, it would be interesting to include measures of key soil enzymes (e.g. cellulase, protease, phosphatase, sulfatase), their potential to release N_2_O and CH_4_, their hydrophobicity and clay effects. A further consideration should be in the sampling design, to ensure equal and adequate representation of soil types in the aggregate vegetation classes in order to accurately capture the interaction effect.

## Supporting information

**Table S2:** Shows conceptually comparable classification of the soils in the World reference base (WRB) Classification. Number in brackets indicates the number of samples for that soil type

**Table S3:** Land class classification with the corresponding land uses

**Table S4:** The cross tabulation table of Aggregate Vegetation classes (AVCs) versus soil types.

## Acknowledgements

We would like to thank Ed Rowe Robert Mills, Robert Griffiths, and David Cooper at the UK Centre for Ecology and Hydrology for help with the experiments and data handling. Countryside Survey (CS) is funded by a consortium of UK government funded bodies led by the Natural Environment Research Council (NERC) and the Department for Environment, Food and Rural Affairs (Defra), which includes the Centre for Ecology and Hydrology, Countryside Council for Wales, Forestry Commission, Natural England, the Northern Ireland Environment Agency, the Scottish Government, Scottish Natural Heritage, and the Welsh Assembly Government. The completion of CS was only possible with the support and advice of many dedicated individuals from these and other organizations. The authors would like to thank the land owners, farmers, and other land managers who gave permission for the field surveyors to collect data and samples from their land; without such cooperation scientific field studies like CS would not be possible. Additionally, the authors would especially like to acknowledge the major contribution made by the team of laboratory staff who processed all the soils and undertook the chemical analyses.

---

**Figure S1.**
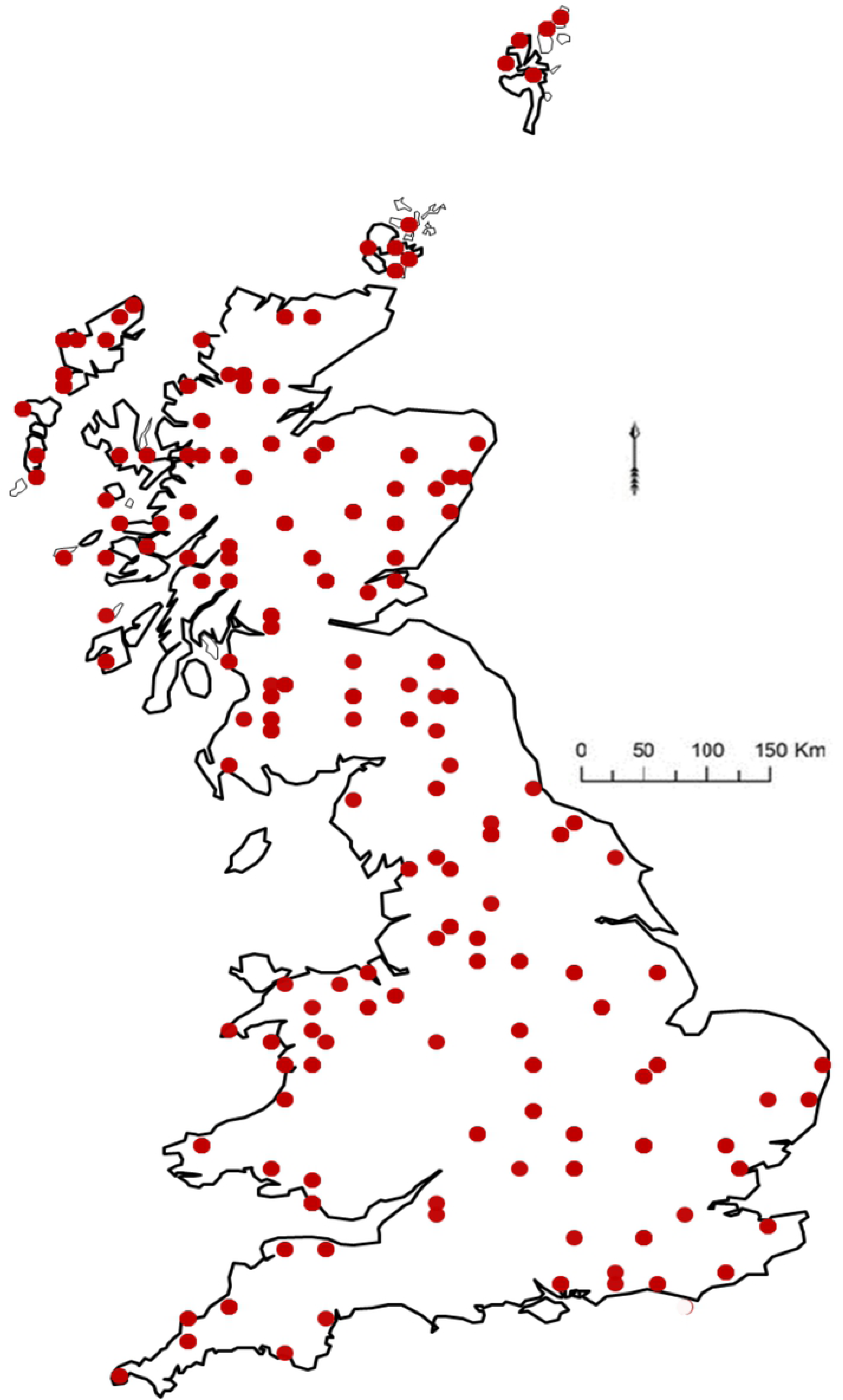
Map of the UK showing the individual soil sampling locations used in the study. The total land area is 209,331 km2.

